# Individuality and information content of infrared molecular profiles: insights from a large longitudinal health-profiling study

**DOI:** 10.64898/2026.04.09.717448

**Authors:** Zita I. Zarándy, Flóra B. Németh, Tarek Eissa, Csilla Lakatos, Dóra Nagy, Diána Debreceni, Frank Fleischmann, Zoltán Kovács, Domokos Gerő, Mihaela Žigman, Ferenc Krausz, Kosmas V. Kepesidis

## Abstract

In this study, we investigate the individuality and information content of infrared molecular profiles derived from blood samples in a large, longitudinal health-profiling cohort and compare them to a standard clinical laboratory panel. Using Fourier-transform infrared spectroscopy, we obtained comprehensive molecular fingerprints from 4,704 self-reported healthy individuals over five visits spanning 1.5 years, alongside routine clinical laboratory measurements. We show that infrared profiles are highly individual-specific and remarkably stable over time, with intra-individual variability significantly lower than inter-individual differences—paralleling the characteristics observed in clinical laboratory data. To quantify and compare the information content of these molecular datasets, we employ individual identification as a proxy for Shannon entropy. In this framework, higher identification accuracy reflects a higher amount of information. Infrared profiles outperform the clinical laboratory panel in identifying individuals at scale, suggesting higher intrinsic information content. Furthermore, combining infrared and clinical laboratory data substantially improves identification performance (the identification of less than 3000 individuals by the clinical laboratory panel is boosted to more than 4000 by incorporating the infrared spectroscopic markers), highlighting the value of integrating complementary data modalities. These findings suggest a practical framework, rooted in information theory, for comparing molecular profiling approaches and emphasize the potential of infrared spectroscopy as a complementary tool in personalized medicine.

## I. INTRODUCTION

Longitudinal molecular profiling has emerged as a promising avenue for personalized health monitoring, offering new possibilities for early disease detection and preventive medicine. The convergence of diverse “omics” technologies—including genomics, transcriptomics, proteomics, metabolomics, epigenomics, microbiomics, and exposomics—alongside biosensor data, advanced imaging, and electronic health records, has created an unprecedented pool of health-related information [1–13]. These integrated datasets are deepening our understanding of human biology and hold the potential to transform how we assess disease risk and detect early warning signs.

Despite this progress, translating molecular insights into clinically actionable tools remains challenging. A key requirement is the ability to quantify how sensitively and reliably different data modalities capture physiological states and health perturbations over time. In this context, simple, minimally invasive, and reproducible approaches are of particular interest.

Infrared molecular fingerprinting (IMF) of blood plasma, based on Fourier-transform infrared (FTIR) spectroscopy, offers a complementary perspective within this landscape. By probing the vibrational signatures of biomolecules, IMF provides a comprehensive molecular snapshot of plasma composition, capturing information across major biochemical classes such as proteins, lipids, carbohydrates, and metabolites [14–18]. Previous studies have demonstrated the stability of IMF over time [19, 20] as a prerequisite for characterizing physiological and pathological states, including applications in cancer detection and metabolic profiling [21–33]. However, large-scale, longitudinal investigations systematically evaluating IMF’s stability, individuality, and clinical relevance are still lacking.

In this study, we take a first step in this direction by analyzing IMF data within a large, prospective health-profiling cohort—Health for Hungary – Hungary for Health (H4H)—designed to monitor health trajectories across the Hungarian population over a decade [34]. We focus on a well-characterized subset of 4,704 self-reported healthy individuals at recruitment, each providing blood samples across five visits spanning approximately 1.5 years. Using 23,520 plasma samples, we comprehensively assess the individuality and temporal stability of IMF signatures (Figure 1), benchmark them against routine clinical blood parameters, and explore their integration.

**FIG. 1.**
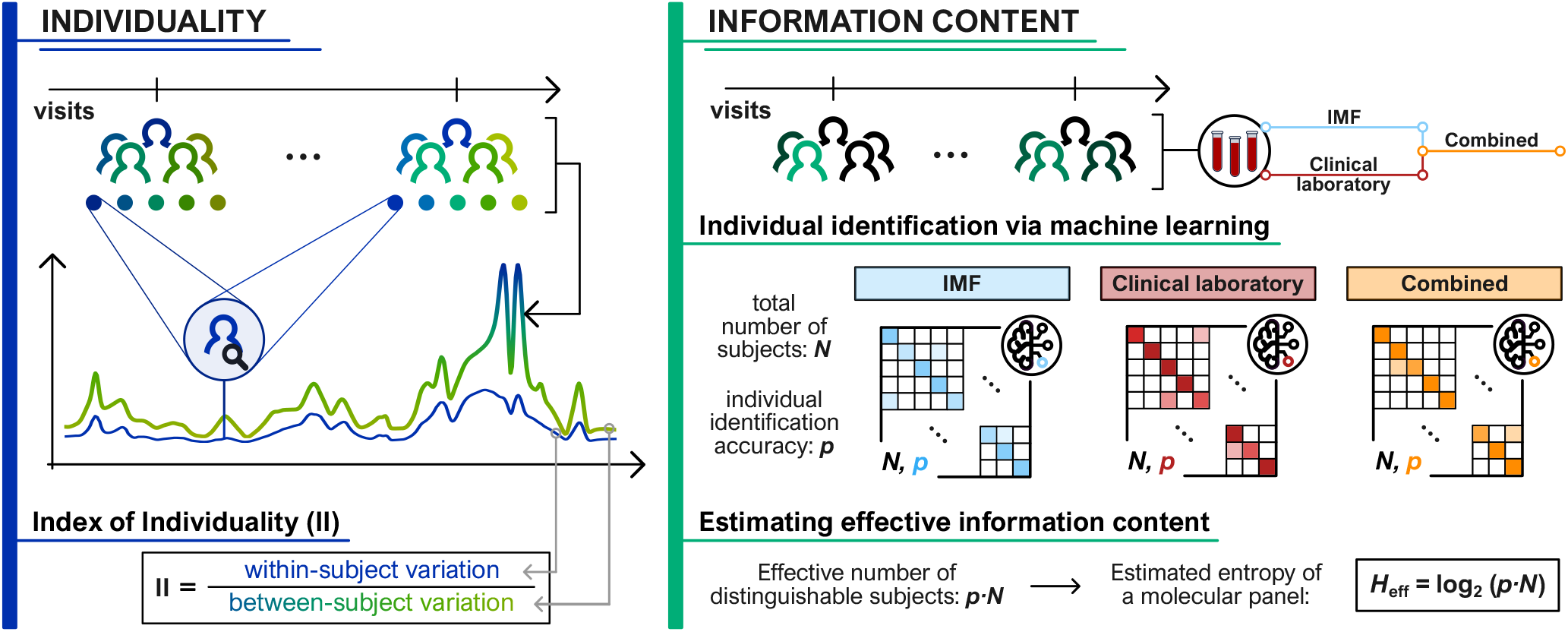
Graphical overview of the study, illustrating its two main focuses: individuality and information content of molecular profiles.

A central aim of our study is also to suggest a framework for quantifying and comparing the information content of different molecular profiling approaches (Figure 1). To this end, we use individual identification as a practical probe for information content, grounded in Shannon’s information theory [35]. In this framework, greater identification accuracy reflects higher information content. Our results show that IMF profiles are not only highly individual-specific and temporally stable but also outperform a standard clinical laboratory panel in large-scale individual identification, suggesting higher intrinsic information content. When IMF and routine clinical laboratory data are combined, identification accuracy improves remarkably, consistent with the information-theoretic principle that entropy increases when combining variables that carry non-redundant information. Moreover, to characterize the shared information content between the mid-IR spectra and the clinical laboratory variables, we performed correlation and regression analyses. Molecules with strong mid-IR absorption, such as glucose (GLU) and lipids, showed high correlation with the measured IMF profiles, and their concentrations could be predicted with comparatively high accuracy using linear regression models. In contrast, enzyme activities such as gamma-glutamyl transferase (GGT) exhibited much weaker correlation with the IMF profiles, and their values were predicted with substantially larger errors.

These findings provide a novel and quantitative approach to assessing the value of molecular profiling technologies, supporting the integration of FTIR-based infrared fingerprinting as a powerful, complementary modality in the future of personalized medicine.

## II. RESULTS

### A. Longitudinal study overview and data modalities

The H4H initiative is a nationwide, prospective cohort study designed to monitor health trajectories across the Hungarian population over 10 years. More than 10,000 participants have been recruited across over 20 study centers, with each individual scheduled for up to 10 follow-up visits. In the present study, we focused on 4,704 participants who completed five study visits, with the first four spaced 130 days apart and the fifth 150 days later (Figure 2a). This yielded 23,520 plasma samples, along-side standardized health questionnaires, clinical laboratory data, and molecular profiling.

**FIG. 2.**
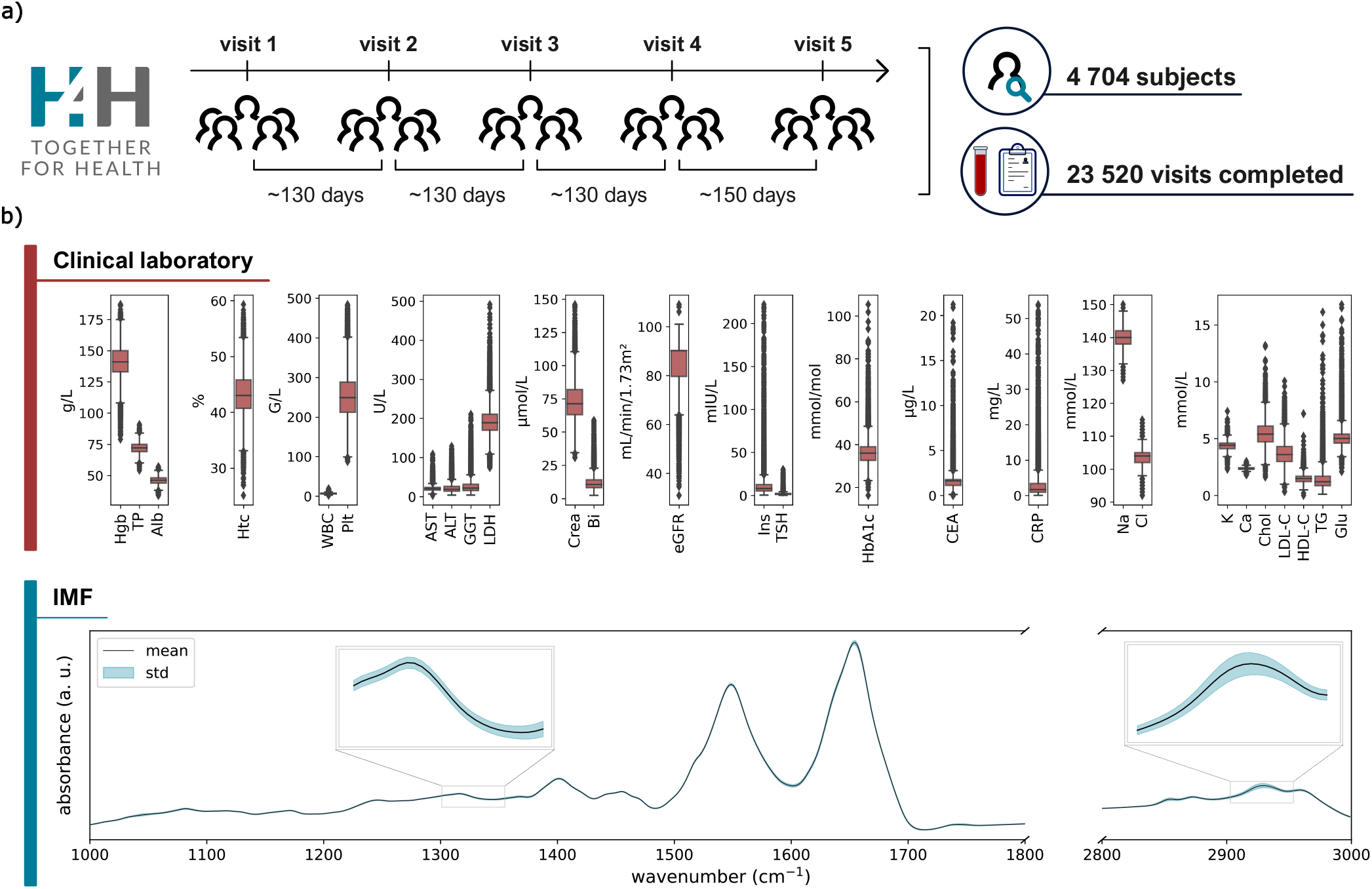
Overview of the H4H (Health for Hungary – Hungary for Health) data and cohort design used in this study. **a)** A total of 4,704 participants each completed five visits over approximately 18 months. At each visit, participants answered questionnaires on demographics, health status, medication use, and comorbidities, and provided a blood sample. **b)** The blood samples were used to generate two data modalities: clinical laboratory measurements and IMF signatures. The figure shows the distribution of 27 clinical biomarkers across all visits and the average absorbance spectrum with standard deviation. All abbreviations for clinical parameters are defined in Appendix Table I.

This study leverages a comprehensive multimodal dataset derived from a longitudinal cohort. Each participant visit includes two distinct data modalities: (i) clinical laboratory measurements encompassing 27 clinical laboratory biomarkers; and (ii) infrared molecular fingerprints obtained from Fourier-transform infrared (FTIR) spectroscopy of plasma samples (Figure 2b). The dataset is accompanied by structured questionnaire responses capturing demographic, lifestyle, and medical history information. All abbreviations for clinical laboratory parameters are defined in Appendix A. Spectral data were acquired using a liquid-phase FTIR spectroscopy under standardized conditions and preprocessed according to established workflows [16, 19, 21, 25, 27, 36]. Together, these modalities provide complementary views into individual health status, enabling the integrated analysis of self-reported, clinical, and molecular data across multiple timepoints. Further details on the data collection, sample handling, demographics and analyses are provided in Appendix B and C.

### B. Individuality of infrared molecular profiles

To assess the suitability of molecular markers for individualized health tracking, we quantified both within-subject and between-subject variation in our two modalities. Within-subject (individual) variation (SD_I_) was calculated as

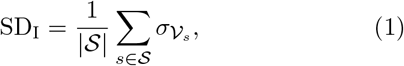

where 𝒮 the set of subjects, 𝒱_*s*_ is the set of repeated visits for subject *s*, and 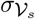 is the standard deviation of the measurements across those visits. Between-subject (group) variation (SD_G_) was analogously determined as

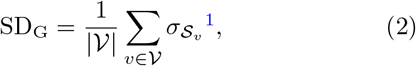

where 𝒱 is the set of repeated visits, 𝒮_*v*_ is the set of subjects measured at visit *v*, and 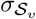 is the standard deviation of the measurements across those subjects.

The Index of Individuality (II) is a widely applied metric in biological variation studies, quantifying how uniquely a biomarker reflects person-specific characteristics relative to population-level variation [37]. The II is defined as the ratio of within-subject to between-subject variation

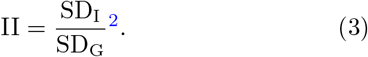

We did not isolate the analytical variation (SD_A_) from SD_G_ and SD_I_ before calculating the II. As a result, both measures include the analytical component, and the reported II values may slightly overestimate the true index. The analytical variation (SD_A_) was derived using repeated quality control measurements, and is shown in Appendix D 1. Reproducibility analysis of the IMF measurements is provided in Appendix D 2.

Low II values (*<* 0.6) indicate that a variable is both stable within a person and sufficiently distinct between individuals—properties desirable for longitudinal monitoring and early disease detection [37, 39]. Biomarkers with higher II values have limited utility for individualized interpretation and are more suitable for population-based reference intervals.

Here we present the first direct comparison of the II between standard clinical laboratory parameters and infrared spectroscopic variables, assessed in a large, well-controlled longitudinal cohort. Using repeated blood samples from 4,704 self-reported healthy participants across five time points, we evaluated II for the 27 clinical laboratory analytes and IMF data.

For clinical laboratory variables, II values spanned a broad range (Figure 3a), reflecting marked differences in their potential for individual-level profiling. Analytes such as CEA and GGT showed the lowest II scores, indicating strong within-person stability and high discriminatory power between individuals, properties that make them promising candidates for longitudinal monitoring. Conversely, electrolytes like sodium, chloride, and potassium exhibited high II values, suggesting limited individual specificity due to high short-term variability. Our results are consistent with the findings reported in the EFLM Biological Variation Database [40], indicating good agreement between the Index of Individuality derived from our cohort and established reference values (Appendix D 3).

**FIG. 3.**
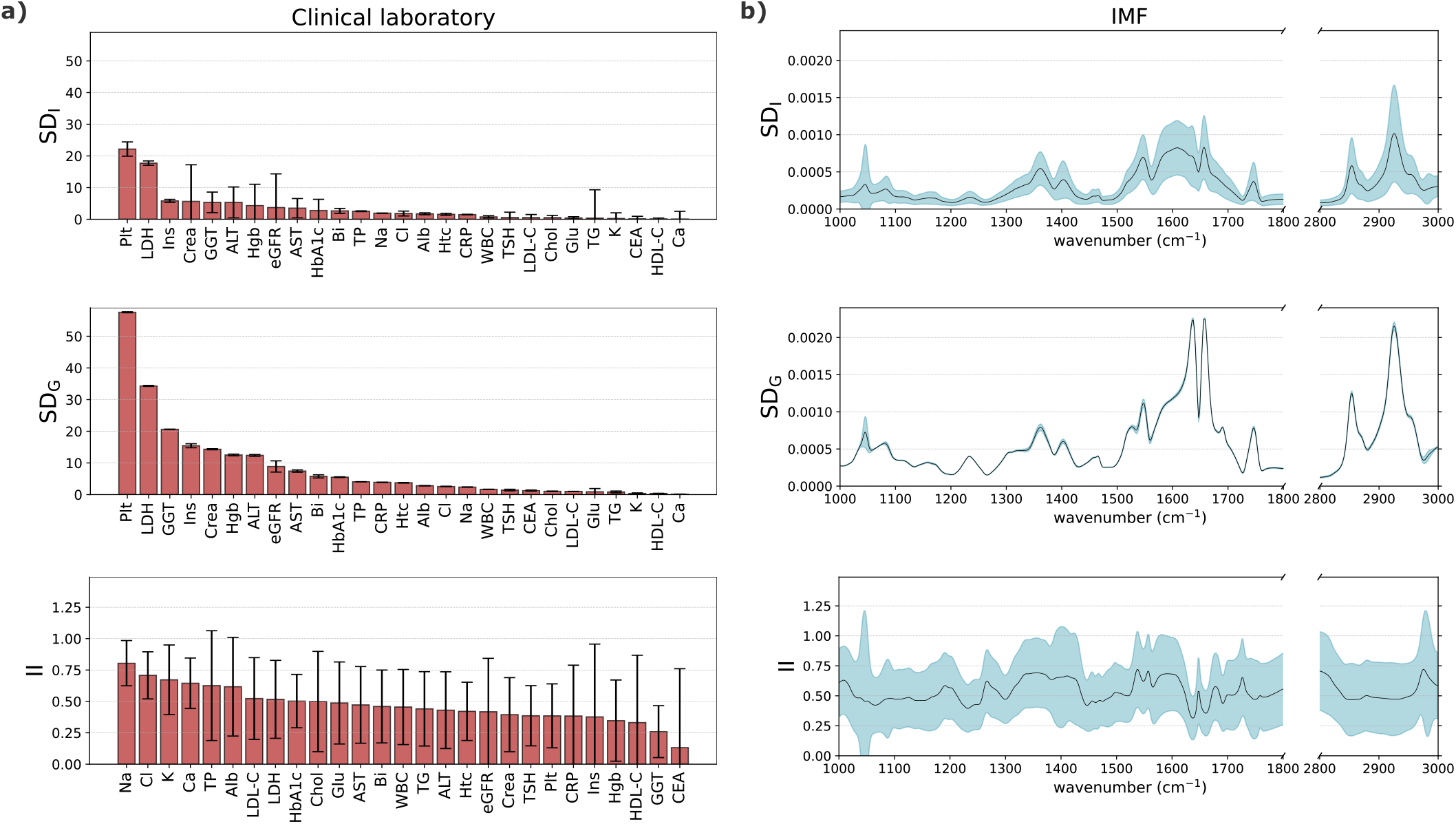
Assessment of variability and individuality across clinical laboratory measurements and FTIR-derived IMF spectra. Three biological variation metrics were evaluated: (1) within-subject variation (SD_I_), estimated across repeated visits for each individual and averaged across subjects; (2) between-subject variation (SD_G_), calculated across individuals at each visit and averaged across visits; and (3) the Index of Individuality, defined as the ratio of within-subject to between-subject variation. **a)** Biological variation of clinical laboratory measurements. **b)** Biological variation of IMF. IMFs were vector normalized using Euclidean *L*_2_ norm before calculating the standard deviations and the Index of Individuality to remove variations in overall intensity arising from differences in sample thickness or path length (see Section B 4 in Appendix).

IMF spectral features displayed a more structured and consistent individuality pattern (Figure 3b). Key biochemical regions—including the CH-stretching (2850–2950 cm^−1^), Amide I (1660 cm^−1^), and Amide II band (1550 cm^−1^)—showed the lowest II values, suggesting they encode stable, person-specific molecular information reflective of proteins, lipids, and other macro-molecular structures [41, 42].

When comparing overall II distributions between modalities (Appendix E), IMF features had a higher average II (0.549 ± 0.090) than clinical laboratory (0.466 ± 0.141), but with lower standard deviation, indicating a more uniformly individual-specific signal across the spectrum. While clinical analytes varied widely in their degree of individuality, IMF provided a consistently information-rich molecular fingerprint, supporting its use as a robust and general-purpose tool for individualized monitoring.

We further examined the robustness of II estimates against potential site-specific biases and found high reproducibility across study sites (Figure 13), validating the reliability of these metrics.

To complement the II analysis, we also assessed intra-class correlation (ICC) as a measure of temporal stability. Variables that combined low II with high ICC were considered especially well-suited for individual tracking. In this regard, CEA emerged as the top-performing clinical marker, with both exceptional stability and person specificity, followed closely by GGT. In the IMF profiles, Amide and CH-stretching regions also demonstrated favorable II–ICC profiles, reinforcing their potential for longitudinal monitoring. In contrast, spectral regions around 1045 and 2980 cm^−1^—associated with increased noise—showed lower stability and higher variability, further validating II as a useful quality filter for biomarker selection (Appendix F).

This first-of-its-kind direct comparison reveals that while selected clinical laboratory analytes offer strong individual-level specificity, FTIR spectroscopy provides a dense and consistent individual signal across the entire molecular spectrum.

### C. Information content and individual identifiability

A central challenge in personalized medicine is comparing the information content of different molecular profiling modalities—particularly when assessing their capacity to capture stable, individual-specific signals from human biosamples. In this work, we introduce and apply a direct, interpretable framework for addressing this challenge by quantifying how well different datasets enable individual identification. This task directly reflects the information content of a molecular profile under Shannon’s information theory [35, 43, 44].

In Shannon’s framework, the entropy of a system measures the average uncertainty associated with a random variable. For a discrete variable with *N* equally probable outcomes, entropy *H* (in bits) is given by

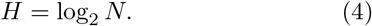

More generally, for a variable with non-uniform probabilities *p*_1_, *p*_2_, …, *p*_*N*_, entropy is defined as

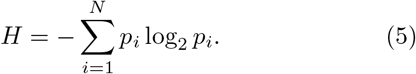

This quantity determines the number of distinguishable states a system can encode. Accordingly, the maximum number of reliably identifiable individual profiles is bounded by

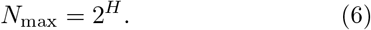

Directly estimating the entropy of high-dimensional molecular datasets is infeasible because of limited sample sizes and the curse of dimensionality [45, 46]. Instead, we use classification performance as an effective probe. We make the simplifying assumption that the number of reliably identifiable individual profiles can be approximated as

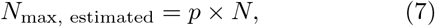

where *p* is the observed multi-class classification accuracy when distinguishing among *N* individuals (classes). Using this, we can derive the lower bound to the Shannon entropy of the dataset in bits from Equation 4, which quantifies the information content associated with correctly identifying an individual from a molecular profile

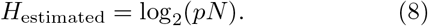

It should be noted that this procedure does not yield the intrinsic entropy of the dataset in isolation. Rather, it characterizes the combined system comprising both the dataset and the employed classification model. As the models were not exhaustively optimized for every cohort size, the resulting estimates reflect a lower bound on the number of individual profiles that a set of biomarkers can support.

Accordingly, a biomarker set with higher entropy encodes a greater amount of individual-specific information, enabling more accurate identification and finer resolution of individual profiles. This framework thus provides a principled and quantitative basis for comparing molecular datasets in terms of their identification capacity—how many distinct individual profiles they can, in principle, distinguish. We characterize this capacity by a practical lower bound derived from observed classification performance log_2_(*pN*).

To operationalize this concept, we trained supervised classification models to identify individuals based on their molecular profiles measured at earlier timepoints. Specifically, we evaluated three feature sets derived from the two previously described modalities: (i) preprocessed IMF spectra, (ii) standard clinical laboratory panel data, and (iii) a combined dataset integrating both modalities. Each model was tasked with predicting individual identity based on longitudinal data collected over five study visits. The experimental design is shown in Figure 4a.

**FIG. 4.**
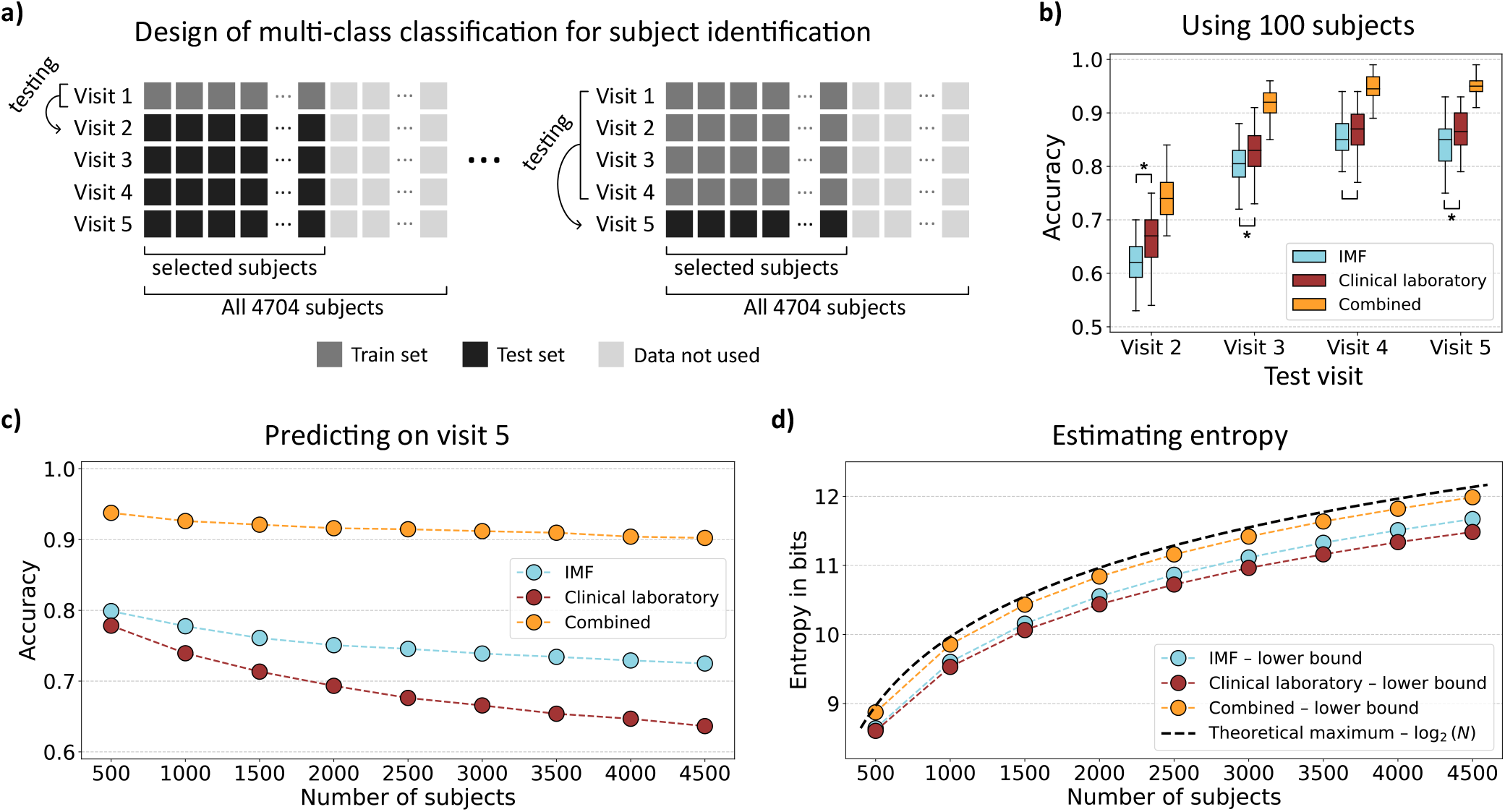
Multi-class classification results for predicting individual identities from longitudinal data across five consecutive visits. **a)** Experimental design: for a randomly selected set of *n* participants, models were trained on the first *i* visits and evaluated on the subsequent one to assess re-identifiability based on prior data. **b)** Learning curves in a fixed-size cohort of 100 participants: classification accuracy was evaluated as a function of the number of visits included in the training set. Each setup was repeated 50 times with different random 100-subject subsets to estimate accuracy distributions. Asterisks indicate statistically significant differences (Mann–Whitney U test), with the combined modality consistently outperforming the individual feature sets. **c)** Scalability analysis: model performance was assessed across increasing numbers of participants, using the first four visits for training and the fifth for testing, to evaluate accuracy decay with cohort size. Each setup was repeated 5 times to gain the mean accuracies presented. **d)** Indirect entropy estimation: for each feature set, the effective number of identifiable individuals was approximated as *p × N* (where *p* is classification accuracy and *N* is cohort size), and converted into an estimated entropy *H*_estimated_ = log_2_(*pN*). This lower bound is compared to the theoretical upper bound for a cohort of *N* individuals, log_2_ *N*, indicated by the continuous black line.

We first evaluated how classification performance improves with increasing temporal data per participant (Figure 4b). For this, we assessed learning curves where classification accuracy was evaluated as a function of the number of visits included in the training set, using the immediate subsequent visit as the test set; for example, if visit *n* is the test, the training set included visit 1 through *n* − 1. In a sub-cohort of 100 individuals, repeated 50 times with different randomization to estimate accuracy distributions, accuracy increased substantially when incorporating data from up to three visits and then plateaued with a fourth. This finding indicates that just three observations are sufficient to capture stable, individual-specific molecular patterns and establish a personalized baseline.

Next, we gradually increased the cohort size *N* used for probing the information content of datasets by individual identification from 500 to 4,500 (Figure 4c). As expected, classification accuracy slightly declined with growing cohort size, reflecting the challenge of resolving more individual profiles. However, models trained on the combined feature set maintained strong performance even at the largest cohort sizes, achieving over 90% accuracy and correctly identifying more than 4,000 of the 4,500 individuals. This confirms that the combined IMF profiles and clinical laboratory data carry sufficient entropy to support very accurate identification and finer resolution of individual profiles across thousands of participants.

Importantly, we observed that across all cohort sizes, the combined modality consistently achieved the highest accuracy, clearly demonstrating that IMF signatures and clinical laboratory data carry complementary information. This aligns with information theory: joint entropy exceeds that of either source only when the added features are not fully redundant—implying that the IMF spectral dataset contains information absent from the clinical panel, and vice versa. By applying our heuristic estimation of effective entropy, we observed a clear separation between the three feature sets, with the combined modality showing the highest *H*_estimated_ across all *N* values, approaching the theoretical upper bound (Figure 4d).

These findings highlight the potential of personal identification analysis not only as a benchmark for the information content of a given set of biomarkers but also as a rigorous tool for evaluating the added value of multimodal integration. For extended performance results (including the exploration of temporal stability), site-specific robustness checks, and detailed modeling configurations, see Appendix G and H.

### D. Interpretation of infrared molecular profiles

Infrared molecular fingerprints encode broad biochemical information reflecting the complex molecular composition of blood plasma. To interpret this signal in a clinical context, previous studies have explored how spectral features relate to routinely measured blood analytes [25]. These investigations revealed that several metabolic markers exhibit strong correlations with specific infrared absorption bands. These band assignments are consistent with previous FTIR plasma studies, including curated biochemical interpretations of absorption features [42].

In this work, we conducted a similar univariate correlation analysis between individual spectral features and 27 clinical blood analytes (Figure 7). Consistent with prior reports, we observed that triglycerides, total cholesterol, and LDL cholesterol showed high *r*^2^ values with IMF values, particularly in the CH-stretching (around 2800–3000 cm^−1^) and C=O-stretching (around 1740 cm^−1^). Glucose, insulin and HbA1c also exhibited distinct *r*^2^ profiles, with all showing concentrated signal around 1000–1150 cm^−1^.

Hematocrit showed a structured and high *r*^2^ profile, and hemoglobin, which is tightly associated physiologically, followed a nearly identical spectral correlation pattern, reflecting their shared erythrocyte origin. Albumin, the most abundant plasma protein, showed strong *r*^2^ signals with the Amide I and Amide II bands (1550–1700 cm^−1^), while total protein showed moderate correlation spanning a broader region (1200–1550 cm^−1^), partially overlapping but distinct from albumin. White blood cells and C-reactive protein (CRP), both markers of inflammation, displayed weaker and more diffuse correlations, without pronounced peaks, suggesting indirect or less specific spectral contributions. These observations support the interpretability of our infrared signals and their relevance to specific molecular classes. However, it should be noted that hemoglobin, hematocrit, white blood cells, and platelets are not plasma-derived markers but are typically measured from whole blood. Therefore, any observed spectral correlations may arise indirectly. The other analytes, including electrolytes and liver enzymes, showed only low or near-zero correlation across the spectrum, indicating limited direct infrared detectability under the current measurement conditions.

To complement these univariate analyses, we applied multivariate regression to predict clinical analyte concentrations from IMF spectra using MultiTask Lasso, which jointly estimates sparse regression models across correlated outputs. Model performance was evaluated using 5-fold cross-validation. Figure 5 summarizes the performance of predicting clinical analytes from FTIR spectra across 27 parameters. Prediction quality is quantified using Pearson correlation coefficients and relative mean squared errors (MSE), aggregated across all cross-validation folds. Error bars represent the standard deviation of performance metrics across cross-validation folds.

**FIG. 5.**
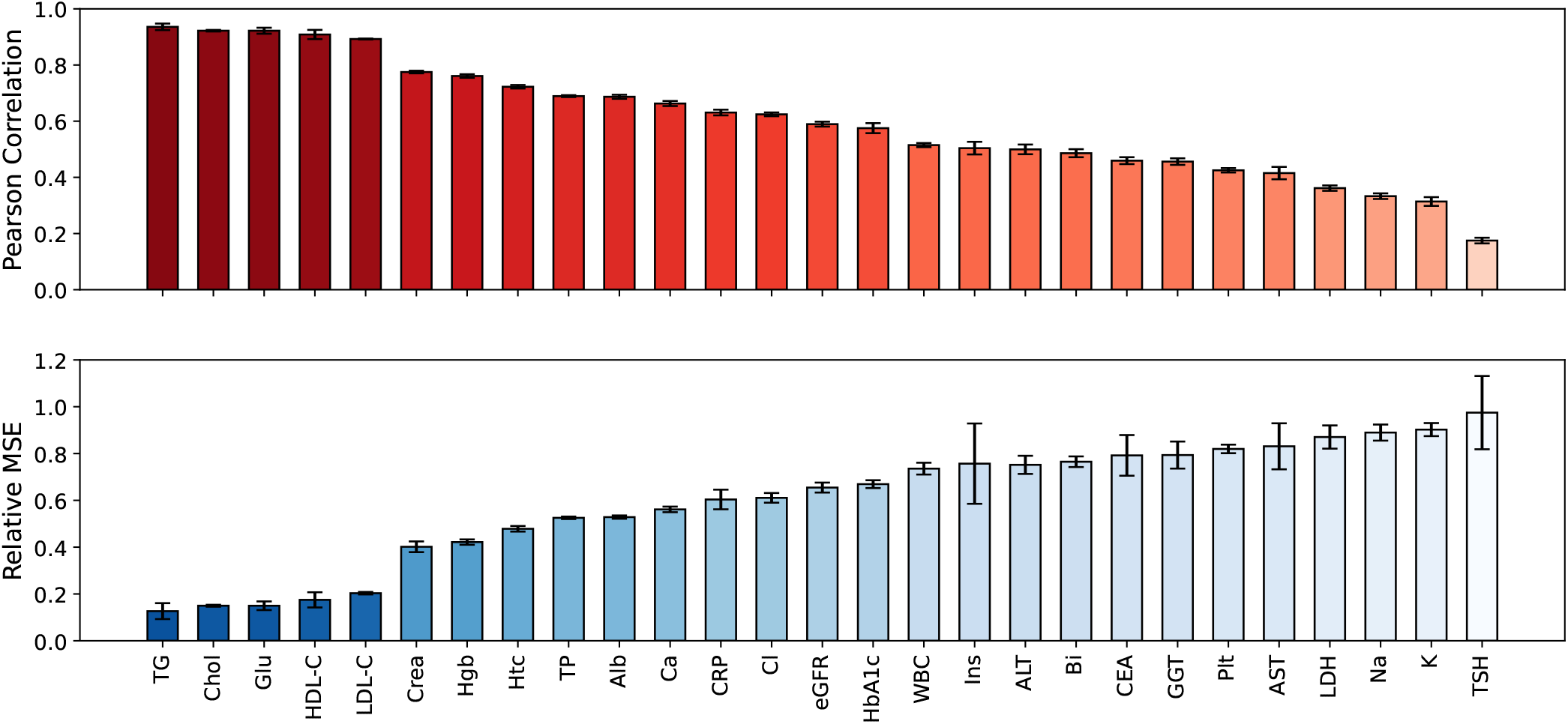
Predictive performance of FTIR spectra for clinical analytes. **a)** Pearson correlation between predicted and measured values across 27 blood parameters. **b)** Relative mean squared error (MSE) of predictions, standardized per analyte. Error bars indicate standard deviation across cross-validation folds.

As expected from the univariate analysis, the top predictive performance was achieved for lipid-related markers and glucose, each reaching Pearson correlations over 0.89. Based on their strong spectral correlation and shared physiological basis, we also anticipated that hemoglobin and hematocrit would exhibit similar regression performance, which was indeed observed. A comparable expectation held for total protein and albumin, given their biochemical relationship and overlapping absorbance regions, though in this case, total protein showed slightly better predictive accuracy.

Interestingly, creatinine, despite showing relatively weak univariate correlations with IMF features, achieved stronger multivariate predictive performance than one might have expected. This discrepancy underscores the value of multivariate modeling, which can extract meaningful signals distributed across the full spectral range, even when no single spectral feature shows a dominant correlation. It highlights the capacity of these models to detect subtle, distributed biochemical patterns not captured by individual feature associations alone.

Overall, these findings confirm and extend earlier work [25] by demonstrating that IMF spectra encode robust and clinically meaningful biochemical information. They highlight the potential of IMF data as a non-invasive surrogate for multiple blood biomarkers, now confirmed in a larger and demographically diverse longitudinal cohort.

## III. DISCUSSION

This study provides compelling evidence that IMF data based on FTIR spectroscopy captures dense, individual-specific, and temporally stable molecular information from blood plasma. By leveraging a large temporally resolved dataset—4,704 individuals followed across five study visits—we demonstrate that IMF signatures show low within-subject variability and high between-subject distinctiveness. These findings confirm and substantially extend earlier observations on IMF stability and individuality [18, 19], now supported by a larger and more heterogeneous population.

A key result of this work is the systematic comparison of IMF profiles with routine clinical laboratory data using an information-theoretic framework grounded in individual identifiability. The notion of using classification accuracy as an effective probe for information content enables direct, interpretable comparisons across diverse molecular modalities. IMF profiles not only matched but significantly exceeded the clinical laboratory panel in their capacity to uniquely identify individuals, indicating a higher intrinsic information capacity. Importantly, combining the IMF with clinical laboratory data substantially improved individual identification, particularly at scale, underscoring the complementary and significantly non-redundant nature of these two modalities.

This insight has practical implications. In large, population-scale cohorts, the ability to resolve individual molecular trajectories is essential for early detection, longitudinal monitoring, and personalized intervention. The demonstrated gain from multimodal integration supports the broader vision of precision medicine, in which diverse biochemical signatures are synthesized to provide a holistic, individual-level view of health.

The individuality analysis based on the II further reinforces the suitability of IMF for personal health monitoring. Spectral features within well-characterized biochemical regions—such as the Amide I/II and CH-stretching bands—showed particularly strong individuality and temporal stability.

To further dissect the relationship between IMF and clinical laboratory data, we complemented our identifiability-based information analysis with multivariate regression modeling. This allowed us to explicitly probe the extent of shared biochemical information between the two modalities. We found that several clinical analytes—most prominently lipid-related markers and glucose—can be predicted from IMF spectra with high accuracy, indicating a clear shared information component.

While our results are promising, several avenues for future development remain. First, IMF’s clinical utility will benefit from technological innovations. Laser-based infrared spectroscopy, especially field-resolved techniques, is advancing the resolution and sensitivity of spectral acquisition [47–49]. These developments may enhance the detectability of low-abundance or weakly absorbing biomolecules, broadening the diagnostic reach of IMF. Second, methodological enhancements—such as information-optimal feature selection and adaptive sampling—could improve signal-to-noise ratios and reduce measurement overhead in large-scale settings [50]. These approaches offer a pathway toward more efficient and scalable deployment of IMF in routine health assessments.

However, several important limitations need to be acknowledged. Our analysis was restricted to self-reported healthy individuals; therefore, the sensitivity of IMF to early, preclinical, or overt disease states remains untested in this cohort. Prior studies suggest that IMF is responsive to pathological conditions [21, 25–27], but a rigorous evaluation in diverse clinical contexts and populations is needed to determine this. Furthermore, while the IMF captures global biochemical composition of blood plasma, its interpretability at the level of individual molecules and specific molecular pathways is currently limited [18]. Integrating IMF with partially orthogonal omics modalities—such as proteomics, metabolomics or lipidomics—may offer deeper mechanistic insights and enhance biological interpretability.

In summary, our findings position IMF of blood plasma as a robust, information-rich, and individual-specific molecular profiling tool that complements standard clinical laboratory testing. Its high temporal stability and capacity for accurate individual identification suggest strong potential for longitudinal health monitoring. With continued technical innovation and clinical validation, IMF may play a pivotal role in the emerging landscape of personalized and preventive medicine.

## ACKNOWLEDGMENTS

We thank all members of the Center for Molecular Fingerprinting (CMF) for fostering a research environment that made this work possible. We are especially grateful to Kamilla Mileant and Ágnes Majárné Kovács for their support with the FTIR measurements, to Krisztián Borbély for his help with data curation, and to Lea Gigou for insightful discussions. We also thank the BIRD team at LMU Munich for their assistance in establishing the facilities and developing the experimental workflows and paradigms for FTIR data acquisition.

This study was supported by the Center for Molecular Fingerprinting Research Nonprofit LLC (CMF), the Frontiers Foundations, the Centre for Advanced Laser Applications (CALA) at LMU Munich, and the Max Planck Institute of Quantum Optics (MPQ). This work is part of Project No. 2020-2.1.1-ED-2022-00213 that has been implemented with the support provided by the Ministry of Culture and Innovation of Hungary from the National Research, Development and Innovation Fund, financed under the 2020-2.1.1-ED funding scheme. ZIZ and FBN were supported by the 2025-2.1.2-EKÖP-KDP University Research Scholarship Programme of the Ministry for Culture and Innovation from the source of the National Research, Development and Innovation Fund.

## DATA AVAILABILITY

Due to privacy regulations under the General Data Protection Regulation (EU) 2016/679, the datasets analyzed in this study cannot be made publicly available. The code used in this study along with realistic simulated data can be accessed at: https://github.com/Data-Science-Group-Attoworld/H4H-individuality-identifiability.

## Appendix A: Abbreviations

**TABLE I.** List of clinical biomarkers and their abbreviations used in the study.

| **Biomarker** | **Abbreviation** |
| --- | --- |
| Hemoglobin | Hgb |
| Hematocrit | Htc |
| White blood cells | WBC |
| Platelet count | Plt |
| Total protein | TP |
| Aspartate aminotransferase | AST |
| Alanine aminotransferase | ALT |
| Gamma-glutamyl transferase | GGT |
| Lactate dehydrogenase | LDH |
| Albumin | Alb |
| Sodium | Na |
| Potassium | K |
| Calcium | Ca |
| Chloride | Cl |
| Total bilirubin | Bi |
| Total cholesterol | Chol |
| Low-density lipoprotein cholesterol | LDL-C |
| High-density lipoprotein cholesterol | HDL-C |
| Triglycerides | TG |
| Creatinine | Crea |
| Glucose | Glu |
| Insulin | Ins |
| Hemoglobin A1c | HbA1c |
| Thyroid-stimulating hormone | TSH |
| C-reactive protein | CRP |
| Carcinoembryonic antigen | CEA |
| Estimated glomerular filtration rate | eGFR |

## Appendix B: Sample and data overview

### 1. Sample collection

This study drew on data from a large-scale longitudinal cohort consisting of participants who self-reported as healthy at the time of recruitment (study code: H4H HU 2020 Sample Collection; ethics approval reference: 2754-11/2020/EÜIG). As part of the study protocol, certified external laboratories measured 27 routine blood parameters, and Fourier-transform infrared (FTIR) spectra were acquired at the laboratory of the Center of Molecular Fingerprinting in Szeged, Hungary. For this analysis, we focused on a subset of 4,704 participants who underwent FTIR measurements exclusively measured with the same device, each with data collected across five visits.

### 2. Sample handling and FTIR measurements

As part of this study, EDTA plasma samples were collected at 20 study centers across Hungary, following harmonized protocols for blood collection and processing. The resulting plasma was divided into 8–10 aliquots of ml each and stored at –80 °C. A second aliquotation step was performed centrally in Szeged, Hungary: one 0.5 ml aliquot per sample was thawed, gently mixed for 30 seconds, and centrifuged. From this, four 90 *µ*l measurement aliquots were prepared and subsequently refrozen at –80 °C. For the FTIR measurements, these aliquots were thawed again, resulting in two freeze–thaw cycles before FTIR analysis.

All FTIR measurements were conducted in a fully randomized and blinded manner. The operators performing the measurements had no access to clinical information or sample identities. A commercial FTIR instrument specifically designed for liquid samples in transmission mode was used for all analyses (MIRA Analyzer, version MA7; CLADE GmbH, formerly Micro-Biolytics GmbH).

Spectra were recorded using a flow-through transmission cuvette with CaF_2_ windows and an 8 *µ*m optical path length. Data were acquired at a resolution of 4 cm^−1^ across a spectral range of 930–3050 cm^−1^. A water reference spectrum was automatically collected after each sample to reconstruct absorbance spectra. Although the manufacturer’s algorithm for this correction is not disclosed, it is known to overcompensate for highly concentrated samples like human plasma. The effective path length was also determined automatically and used for spectral adjustment. The MIRA-Analyzer does not provide access to raw measurements but only to the water-compensated absorbance spectra.

Samples were processed in batches of 28, with DMSO_2_, water, and pooled human serum (BioWest, Nuaillé, France) included as quality controls, yielding a total of 34 samples per batch. The BioWest quality control samples enabled tracking of analytical variation and instrument stability over time.

### 3. Cohort and demographics

The cohort consists of 4,704 participants, with a broad age distribution centered around 50 years. Body mass index (BMI) shows moderate variability, skewing toward the overweight range. The sex distribution is imbalanced, with a higher proportion of female participants. The age and BMI distributions of the cohort at inclusion, stratified by sex, are shown in Figure 6. These visualizations are complemented by the detailed sample counts in Table II, which summarize the distribution of collected blood samples across age and BMI categories. The cohort includes a higher proportion of female samples. Age distributions are centered around 50 years for both sexes. BMI values span a broad range, with male samples showing slightly higher median values than female samples.

**FIG. 6.**
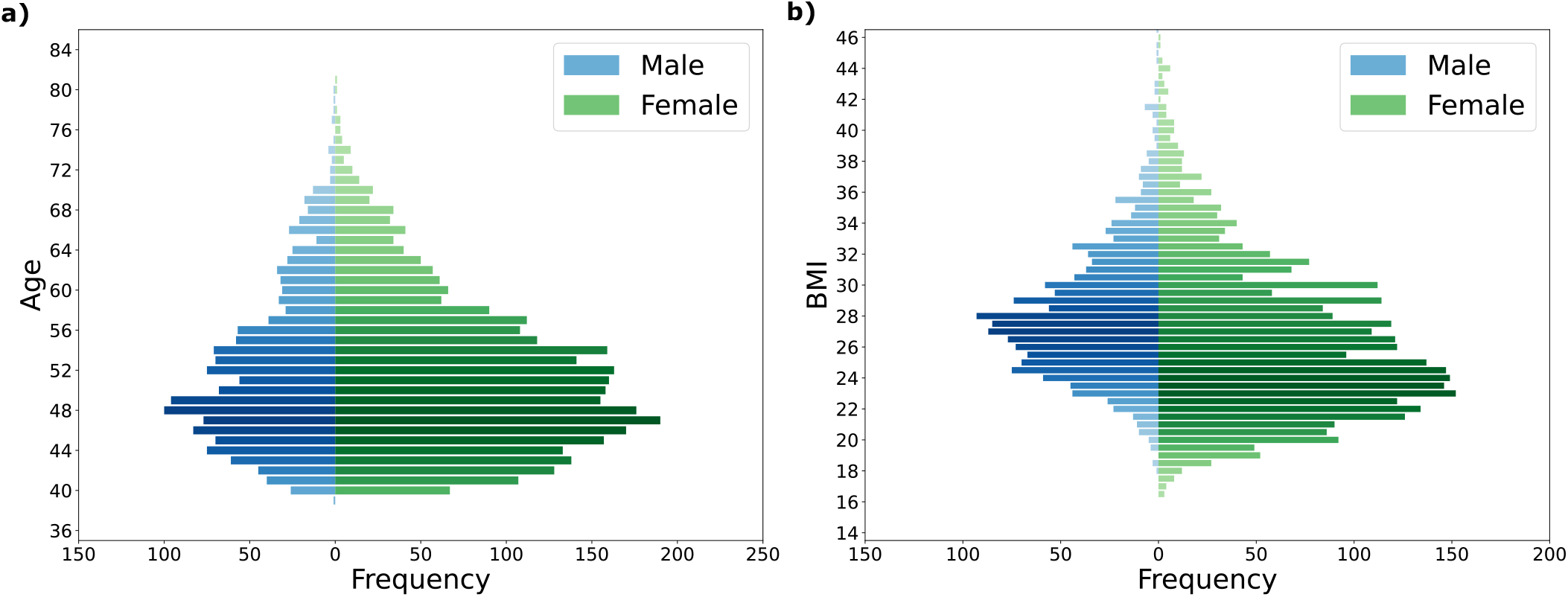
Demographic distribution of the analyzed cohort by sex at inclusion. **a)** Age distribution shows a balanced representation across sexes, centered around 50 years, with a higher number of female participants. **b)** BMI distribution indicates a broad range for both sexes, with males exhibiting a higher median BMI compared to females.

**TABLE II.** Demographics of study participants.

| Category | Range | Male Samples | Female Samples | Total Samples |
| --- | --- | --- | --- | --- |
| Age (years) | 30–39 | 1 | 0 | 1 |
|  | 40–49 | 673 | 1421 | 2094 |
|  | 50–59 | 556 | 1271 | 1827 |
|  | 60–69 | 243 | 435 | 678 |
|  | 70–79 | 30 | 71 | 101 |
|  | 80–89 | 1 | 2 | 3 |
| BMI | 0–18.4 | 1 | 27 | 28 |
|  | 18.5–24.9 | 318 | 1372 | 1690 |
|  | 25.0–29.9 | 735 | 1049 | 1784 |
|  | 30.0+ | 450 | 752 | 1202 |
| **Total** | – | 1504 | 3200 | 4704 |

Molecular profiles were acquired from plasma samples at each visit using Fourier-transform infrared spectroscopy. Additionally, 27 clinical biomarkers were measured through routine laboratory tests, encompassing a wide range of physiological domains, including lipid metabolism, kidney function, inflammation, and hematological indices.

### 4. Data preprocessing

Raw IMF spectra were preprocessed using a four-step protocol to retain informative signal regions and prepare the data for machine learning. First, spectra were truncated at 1000 and 3000 cm^−1^ to remove noisy regions caused by the low transmittance of CaF_2_ below 950 cm^−1^ and the high water absorption above 3000 cm^−1^. Second, a two-stage outlier removal process was applied. Spectra exhibiting an additional, atypical absorption feature around 1045 cm^−1^—presumably resulting from a recurring but uncharacterized source of sample contamination—were excluded. Subsequently, statistical outliers were identified using the Local Outlier Factor (LOF) algorithm applied to the truncated spectra. Outlier filtering removed approximately 6% of samples from the IMF data. Only inlier spectra were retained for further processing. In the third step, spectra were vector normalized using the Euclidean (*L*_2_) norm. This procedure preserves the relative shape of the spectra, including peak positions and ratios, while eliminating effects of absolute scaling. The resulting normalized spectra are therefore directly comparable across samples and suitable for multivariate analyses, ensuring that differences reflect underlying compositional variation rather than measurement artifacts. Finally, the uninformative region between 1800–2800 cm^−1^ was removed. The retained wavenumber ranges for analysis were 1000–1800 cm^−1^ and 2800–3000 cm^−1^, yielding a final feature set of 519 spectral points.

To assess the effectiveness of the preprocessing protocol, classification performance was evaluated in a repeated multiclass setting, where models were trained to identify 100 individuals from their first four visits and tested on the fifth visit, with subject subsets resampled across 100 runs (III). Excluding the spectrally uninformative “silent” region (1800–2800 cm^−1^) consistently improved model performance, as shown by higher accuracies for the no silent region datasets compared to the corresponding full spectra (e.g., *µ*_Acc_ = 0.845 for processed: no silent region vs. 0.807 for processed: full spectra, and 0.679 vs. 0.625 in the raw case). This indicates that removing regions dominated by noise or weak signal enhances the discriminative power of the models in this identification task. Notably, the only silent region datasets still achieved accuracies well above random chance (e.g., *µ*_Acc_ = 0.211 and 0.313), suggesting that this region contains some subject-specific information. However, combining it with the more informative spectral regions did not improve—and in fact reduced—overall performance, implying that the additional features introduce variability that outweighs their limited signal contribution. This pattern is consistent across both raw and processed data, confirming the robustness of the effect.

**TABLE III.** Model performance across different feature sets.

| Dataset | $N_{\text{feat}}$ | $\mu_{\text{Acc}}$ | $\sigma_{\text{Acc}}$ |
| --- | --- | --- | --- |
| raw: full spectra | 1037 | 0.625 | 0.037 |
| raw: no silent region | 519 | 0.679 | 0.042 |
| raw: only silent region | 518 | 0.211 | 0.039 |
| normalized: full spectra | 1037 | 0.807 | 0.035 |
| normalized: no silent region | 519 | 0.845 | 0.034 |
| normalized: only silent region | 518 | 0.313 | 0.045 |

Furthermore, preprocessing (in particular, vector normalization) led to systematically higher accuracies across all feature sets (e.g., processed: full spectra vs. raw: full spectra), supporting its use to improve comparability between spectra and to mitigate intensity-related artifacts.

Clinical laboratory data were preprocessed through unit standardization and outlier removal. Biological outliers were filtered using predefined upper and lower physiological limits for each parameter. Additionally, DBSCAN-based clustering was applied individually to each marker to detect statistical outliers, which were then set to missing values. The proportion of excluded values was substantially lower than for IMF, ranging from 0.005% for cholesterol-related analytes to 0.8% for insulin. Any sample with at least one missing data point after preprocessing was excluded, resulting in a final dataset comprising 4,704 subjects.

### 5. Machine learning analysis

All machine learning analyses were conducted using the scikit-learn library [51] in Python. Before modeling, both the IMF spectral features and clinical laboratory variables were standardized using z-score normalization (StandardScaler), ensuring that each feature had zero mean and unit variance. To prevent data leakage, standardization was always performed after the train-test split. Specifically, the scaler was fit using only the training data, and the learned parameters (mean and standard deviation) were then applied to transform both the training and test sets. When cross-validation was used, standardization was performed independently within each fold: for each training fold, a new scaler was fit on that fold’s training data and applied to both the training and validation data within the fold. All modeling steps, including regression, classification, and cross-validation, were performed on these properly standardized datasets. For the multi-class classification tasks—described in *Information content and individuals identifiability* section—logistic regression models were trained with *L*_2_ regularization, *λ* = 0.1, ‘liblinear’ solver, selected via hyperparameter tuning [51].

## Appendix C: Statistical methods

### 1. Intra-Class Correlation (ICC)

To evaluate the temporal stability of each variable, we computed the Intra-Class Correlation (ICC), which quantifies the proportion of total variance attributable to between-subject differences. ICC was calculated using a two-way mixed effects model, which accounts for fixed measurement occasions and estimates the consistency of individual-level measurements across time

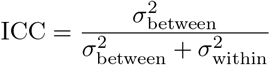

where 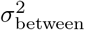 is the variance between individuals and 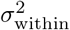 is the residual variance within individuals across visits. ICC values range from 0 (no stability) to 1 (perfect stability), and were computed separately for each clinical laboratory marker and each IMF spectral feature.

### 2. Error propagation for the Index of Individuality

The Index of Individuality (II) was computed as the ratio of within-subject (SD_I_) and between-subject variation (SD_G_). To quantify the uncertainty in II, we applied classical error propagation rules based on a first-order Taylor expansion. Suppose *f* (*x, y*) is a function of two independent random variables *x* and *y* with known standard errors *σ*_*x*_ and *σ*_*y*_. Then the variance of *f* is approximated by:

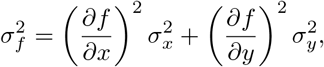

assuming that *x* and *y* are uncorrelated, so their covariance term vanishes. We apply this rule to the function:

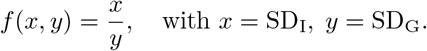

The partial derivatives of *f* with respect to *x* and *y* are:

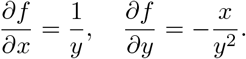

Substituting into the propagation formula gives:

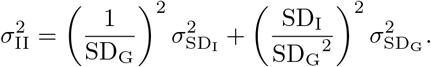

We can factor this to obtain a more interpretable form:

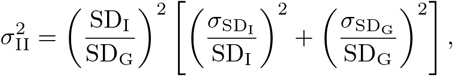

which leads to the final expression:

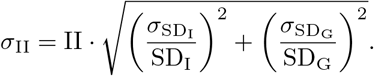

This accounts for the variability in both components while omitting the covariance term.

Assumptions and notes:

- We assume that SD_I_ and SD_G_ are statistically independent, as they are derived from different levels of variability.
- The propagation is based on a linear approximation, which is valid when the uncertainties 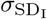 and 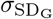 are small relative to the values themselves.
- The computed *σ*_II_ provides an estimate of the standard error of the II, which can be used to plot confidence intervals or quantify uncertainty across features.

This derivation ensures that the reported II values are not only point estimates but are also accompanied by rigorous estimates of their uncertainty, facilitating more robust interpretation and comparison across features or measurement modalities.

### 3. Shared information between clinical chemistry parameters and IMF

Figure 7 shows the univariate squared Pearson correlations between spectral features and clinical analytes, revealing region-specific correlation patterns across lipid, protein, and carbohydrate regions. These *r*^2^ values reflect the proportion of variance in each analyte explained by individual spectral features.

**FIG. 7.**
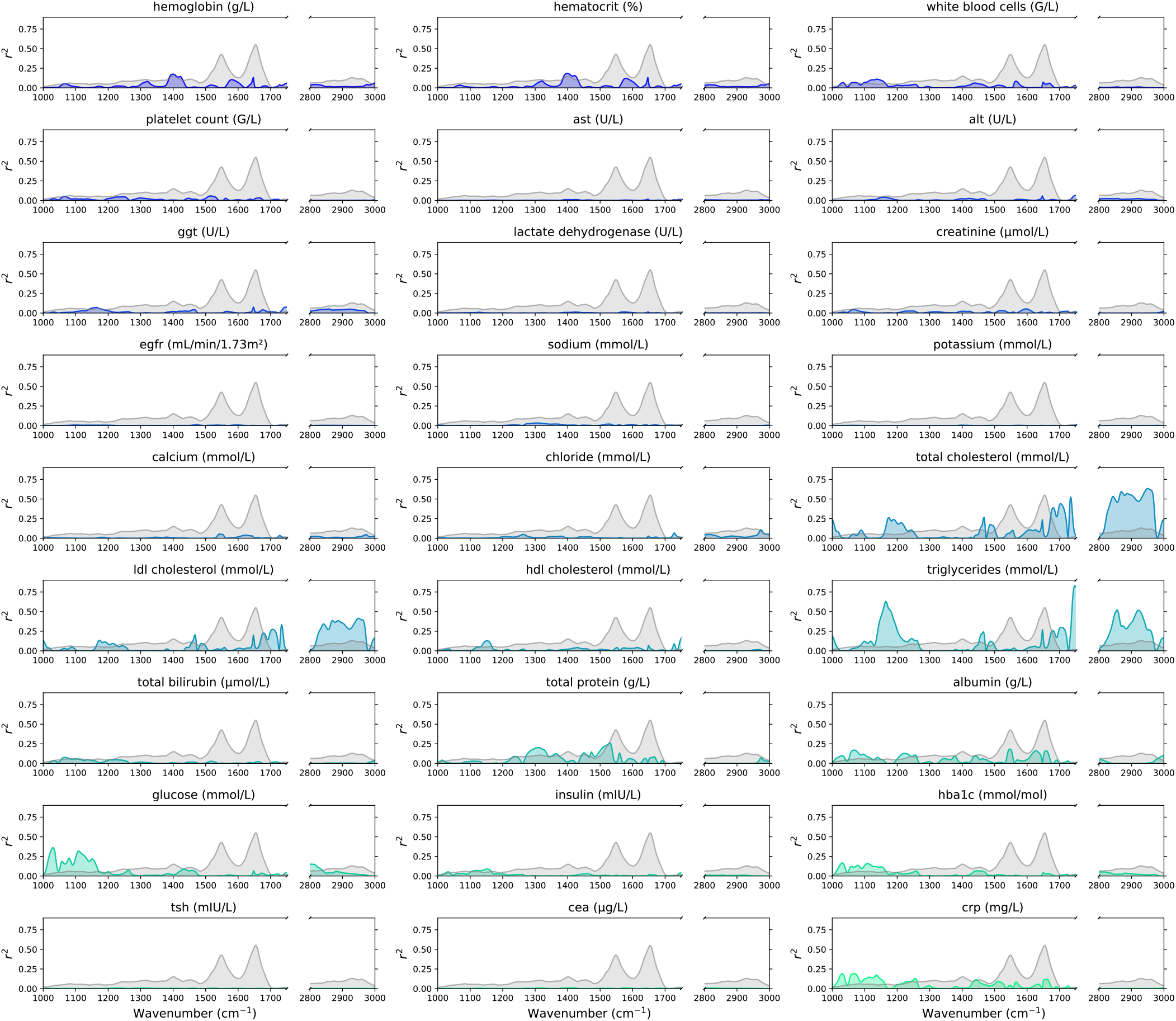
Squared Pearson correlation between each analyte and absorbance values across the 1000–1800 and 2800-3000 cm^−1^ range. The mean IMF spectrum is shown in gray for reference.

## Appendix D: Reproducibility

### 1. Analytical variation of the IMF spectra

For the FTIR instrument, repeated quality control measurements were available, enabling the estimation of analytical variation as the standard deviation of these measurements. To ensure that this estimate is representative of the timescale of the study, the average duration between the first and last FTIR measurement per individual was calculated, yielding approximately 140 days. Based on this interval, the quality control data were divided into three consecutive 140-day periods. Standard deviations were calculated for each period, and the mean and associated uncertainty across the three periods were then used to derive a robust estimate of analytical variation (Figure 8a). The analytical variation closely approaches the within-subject variation in certain spectral regions where the blood sample contains little or no biological information. The so-called “silent region” begins around 1750 cm^−1^, beyond this point, the analytical variation exceeds the within-subject variation, which is consistent with the absence of a meaningful biological signal in this region [21].

**FIG. 8.**
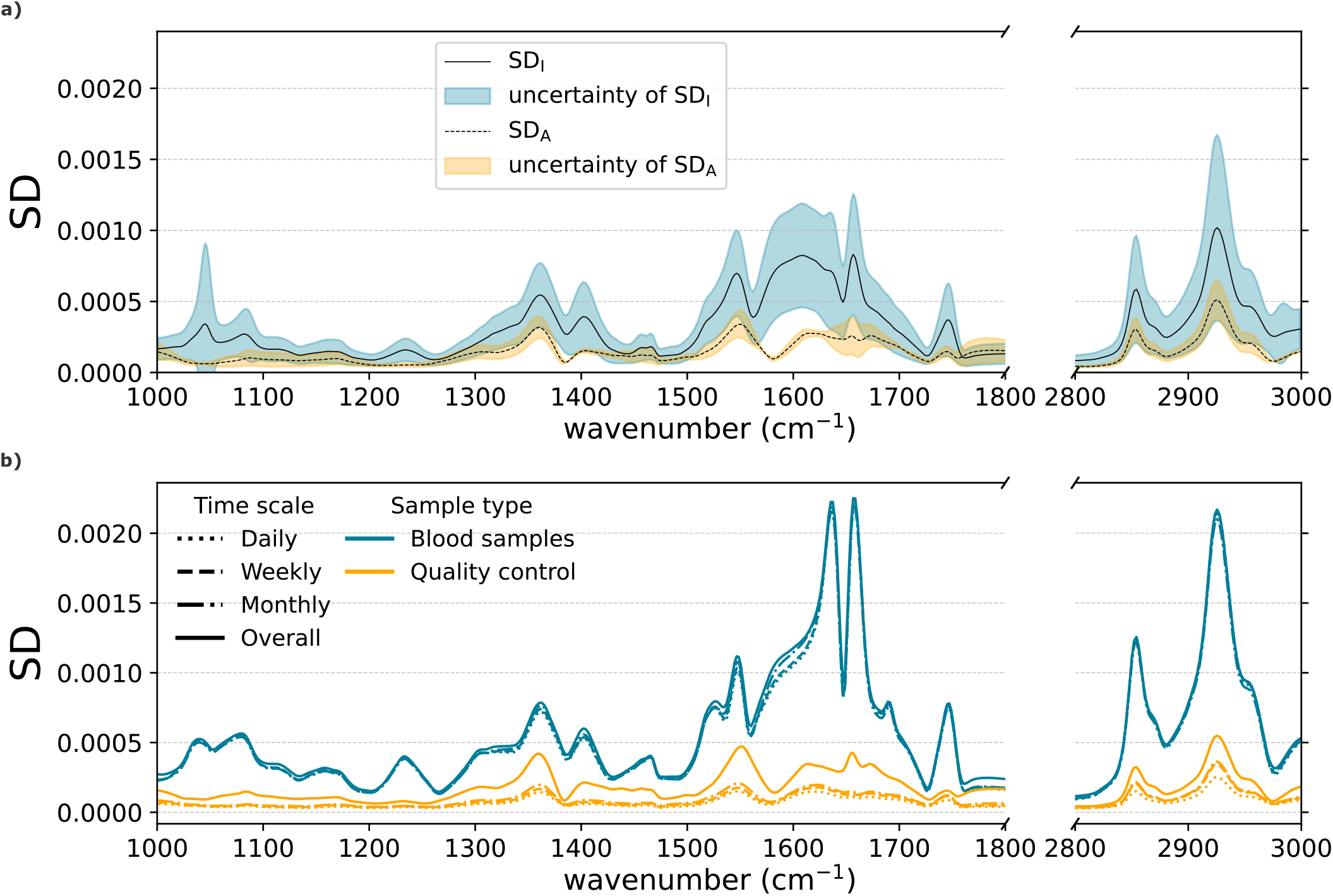
Comparison of blood sample and quality control variability of the IMF spectra. **a)** Within-subject (SD_I_) and analytical variation (SD_A_). **b)** Between-subject and analytical variation across four time scales.

### 2. Quality controls

To assess the reproducibility of our measurements and facilitate comparison with earlier work [48], we followed the same methodology to calculate the wavenumber-averaged standard deviation for both quality control samples and study plasma samples, across predefined temporal groupings. Briefly, measurements were grouped by calendar time (day, week, month), and within each group, the standard deviation was calculated independently at each wavenumber. These per-wavenumber standard deviations were then averaged across the wavenumber dimension, yielding a single variability estimate per time window as shown in Figure 9. The only difference is that, whereas the previous study evaluated the 950–1375 cm^−1^ wavenumber range, we calculated over the 1000–1800 cm^−1^ and 2800–3000 cm^−1^ ranges. To ensure that the standard deviation between individuals is not artificially reduced by the similarity of repeated measurements from the same person, only one sample per individual was retained within each period, selected at random.

**FIG. 9.**
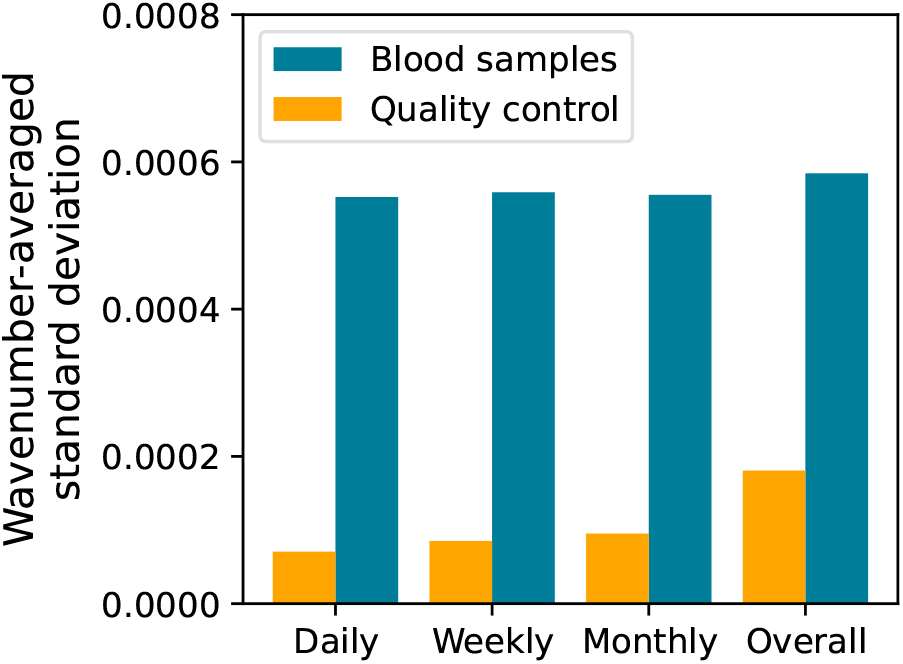
Wavenumber-averaged standard deviation values for quality control samples and study samples across four time scales.

The resulting values are smaller than those reported previously for both quality control and study plasma samples when calculated over the wider wavenumber range. Moreover, the ratio of quality control to study plasma sample variability over the entire campaign is also reduced, which is a desirable outcome. Examining the temporal trends in more detail, we observed that the increase in variability occurs predominantly between the monthly and overall time scales, while differences between shorter intervals remain comparatively small. To better understand the origin of this increase, we subsequently evaluated the variability as a function of wavenumber (Figure 8b) rather than relying on averaged values. This analysis revealed that, in the quality-control samples, the increase in variability is observed across the entire spectral range, indicating a global effect. In contrast, in the analysis of plasma samples, the increase is primarily localized to the amide I and amide II regions, with only minor changes observed elsewhere in the spectrum. Most importantly, these results show that the analytical variability reflected by the quality-control samples remains lower than the biological variability of the plasma samples across the entire spectral region, even over the timescale of the full measurement campaign.

### 3. Comparison to EFLM BV Database

To evaluate the individuality of clinical laboratory parameters in our cohort, we compared the Index of Individuality derived from our data to reference values provided by the EFLM Biological Variation Database [40]. For most parameters, reference values for within-person and between-person biological variation were available and used to compute the corresponding II. However, eGFR is a derived measure rather than a directly assayed analyte, and no biological variation data for it was available in the database.

For each parameter, we used meta-analyzed within-subject and between-subject biological variation estimates published by the EFLM Working Group on Biological Variation, which are derived using standardized BIVAC criteria to ensure high methodological quality. These were then compared to II values calculated from our cohort.

Figure 10 visualizes this comparison. Red bars represent II values from our study, while blue bars represent corresponding EFLM reference values. A notable finding is that the overall trends in II values are consistent with those reported in the literature.

**FIG. 10.**
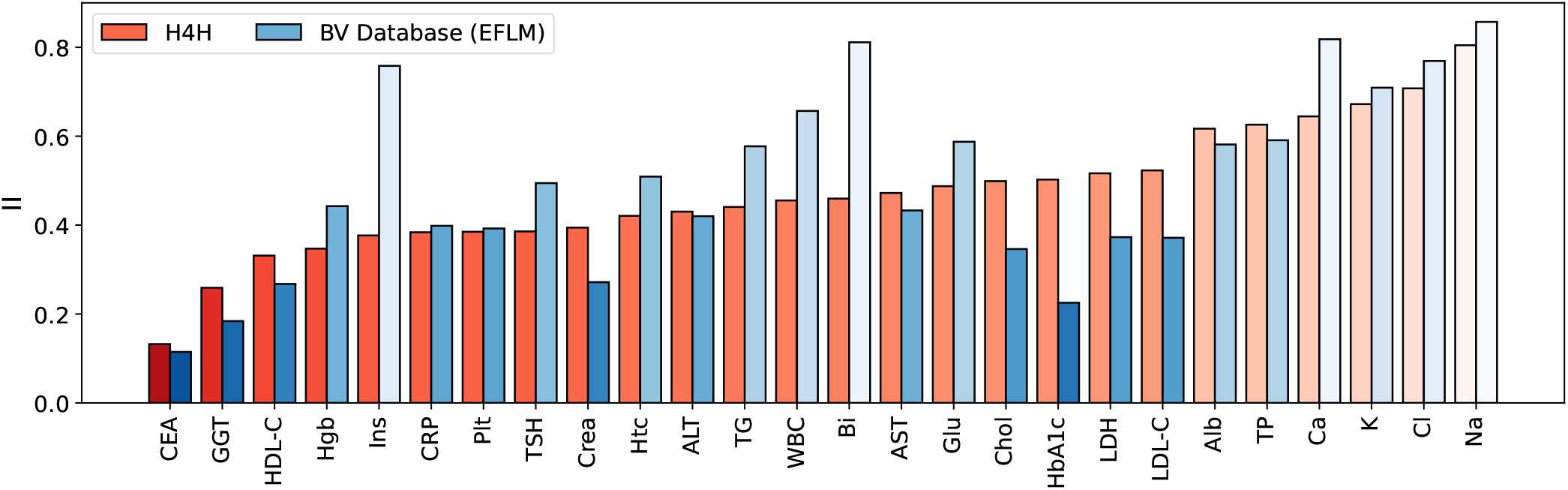
Comparison of Index of Individuality between values derived from the H4H cohort and reference values from the EFLM BV Database.

## Appendix E: Comparison of the II of IMF and clinical laboratory features

When comparing the distributions of II between modalities (Figure 11), IMF features (FTIR; *n* = 519) exhibited a higher mean II (0.549 ± 0.090) than clinical laboratory features (*n* = 27, 0.466 ± 0.141). The IMF values were also more tightly clustered, as reflected by their lower standard deviation, suggesting a more consistent individual-specific signal.

**FIG. 11.**
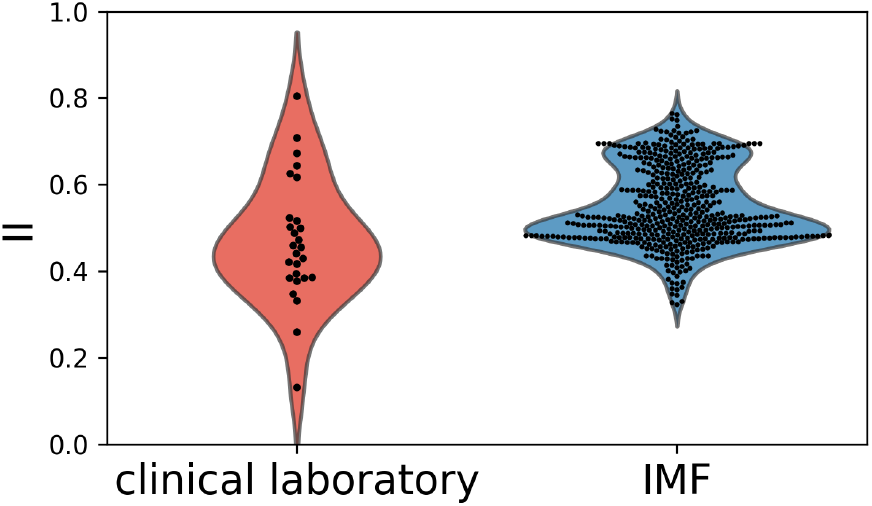
Distribution of II across all features, comparing IMF signatures and clinical laboratory.

## Appendix F: Comparison of II and ICC

To better understand the balance between stability and individuality in our dataset, we examined the Intra-Class Correlation (ICC) alongside the II for both clinical laboratory and FTIR measurements (Figure 12). These two metrics provide complementary perspectives: while II reflects how distinct an individual’s measurements are relative to population variability, ICC quantifies the proportion of total variance attributable to between-subject differences, offering a measure of reliability or temporal stability.

**FIG. 12.**
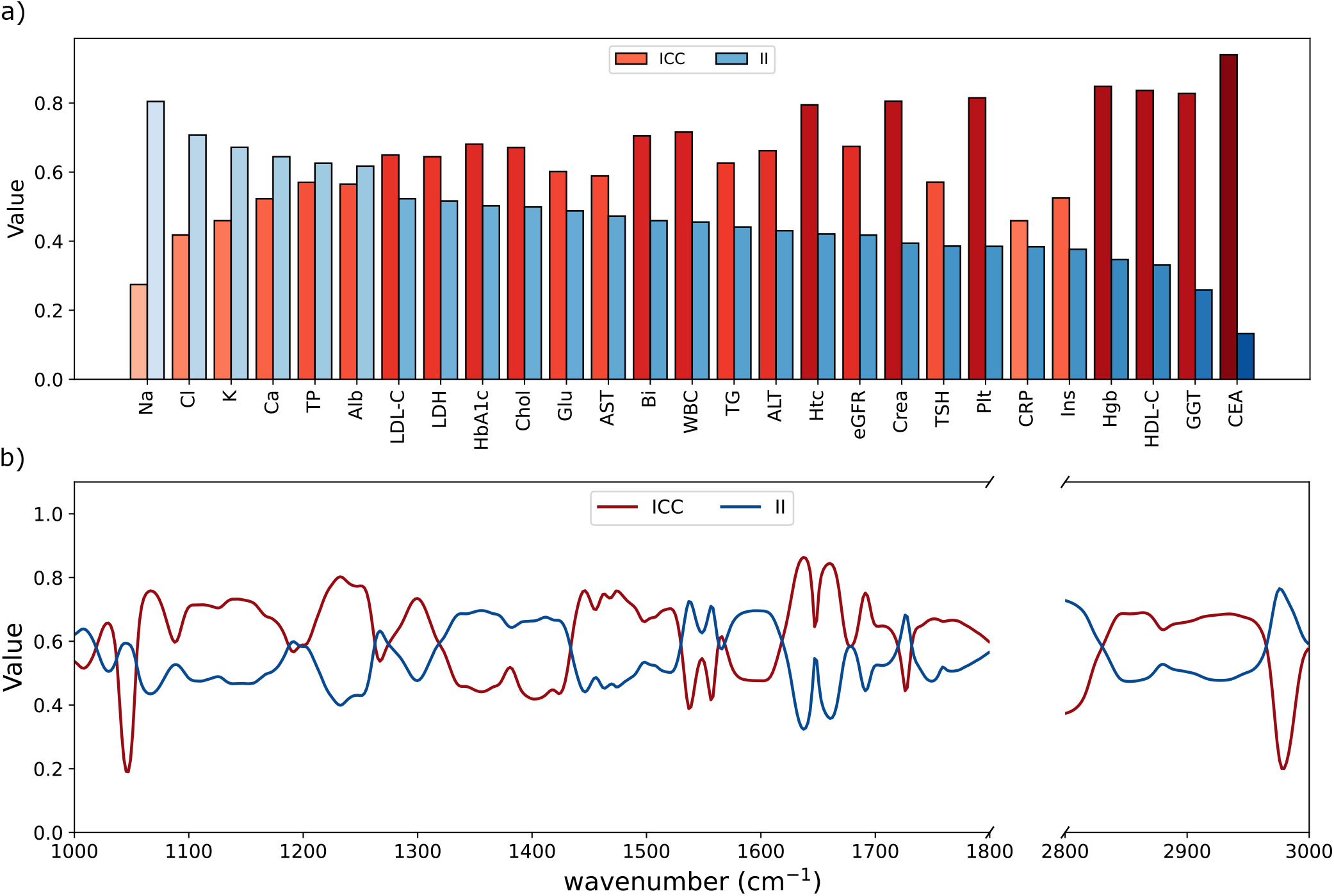
Comparison of Intra-Class Correlation and Index of Individuality across clinical laboratory parameters and the IMF spectral domain. **(a)** ICC and II values for analytes. Darker red bars indicate higher ICC, while darker blue bars indicate lower II. **(b)** ICC and II profiles across the wavenumber range. The ICC represents the stability of each spectral bin, the II reflects spectral individuality. Together, these metrics highlight features suitable for robust and personalized longitudinal monitoring.

Lower II values indicate that an individual’s measurements are relatively stable over time and distinct compared to the broader population—an essential property for personalized interpretation and longitudinal tracking. In contrast, high II values reflect either high within-subject variation or low between-subject variation, reducing the usefulness of the marker for individualized assessments. ICC, in this context, quantifies the proportion of total variance attributable to between-subject differences. Higher ICC values imply stronger consistency of measurements within individuals across time.

Panel (a) displays the ICC and II values of the clinical laboratory parameters. CEA, hemoglobin, HDL cholesterol, and GGT demonstrated a favorable profile of low II and high ICC. These markers are both temporally stable and individually distinctive, making them well-suited for personalized baselines and reliable longitudinal monitoring. On the other hand, CRP and insulin showed low II and low ICC, indicating individualized but dynamic behavior. Their high biological reactivity may limit their use for baseline tracking but make them informative for capturing acute physiological changes.

Some features, including hematocrit and creatinine, exhibited moderate-to-high ICC but relatively higher II, suggesting consistent values over time but insufficient between-person variation for individualized use. These markers may be more informative in group-level or clinical reference ranges rather than in personalized assessments. Finally, markers like sodium and chloride showed high II and low ICC, reflecting poor individual stability and weak population differentiation—traits that reduce their relevance for both longitudinal and personalized profiling.

Panel (b) presents the ICC and II across the spectral range. As with the clinical laboratory data, ICC highlights regions of temporal stability, while II identifies areas where individual spectral patterns are distinct. While the two curves largely mirror each other around a central baseline near 0.6, deviations from this symmetry reveal important insights. In particular, regions around 1040–1070 cm^−1^ and 2960–2985 cm^−1^ show a significant drop in ICC without a comparable rise in II. These ICC dips suggest low temporal stability without corresponding inter-individual variability, which is indicative of measurement noise or an unstable signal rather than biologically meaningful variation. Despite such localized discrepancies, the overall profiles support the conclusion that the Amide and CH-stretching bands are optimal targets for individualized FTIR-based biomolecular profiling, combining both reproducibility and person-specific signal.

## Appendix G: Site-specific bias in clinical laboratory data

To evaluate the potential bias introduced by our simple standardization approach—limited to unit harmonization and the application of StandardScaler from Scikit-learn—we investigated whether site-specific measurement methodologies (causing slightly differing reference intervals reported by laboratories) affected down-stream analyses. No further harmonization was performed to address site-dependent variations in reported normal ranges.

To this end, we calculated the Index of Individuality using subsets of 100 participants matched on age, sex, and BMI, drawn from the five largest study sites (Site A: 590 participants; Site B: 650; Site C: 563; Site D: 844; Site E: 526), as well as 100 matched participants randomly selected from the full cohort of 4,704 individuals. As shown in Figure 13, the II values across the 27 clinical laboratory parameters appeared at a similar level across sites, suggesting minimal site-specific bias. An exception was observed for carcinoembryonic antigen (CEA), due to lower reporting limits at some sites, where values appear capped at 1.63, thereby precluding the quantification of SD_I_ and the SD_G_ is impossible.

**FIG. 13.**
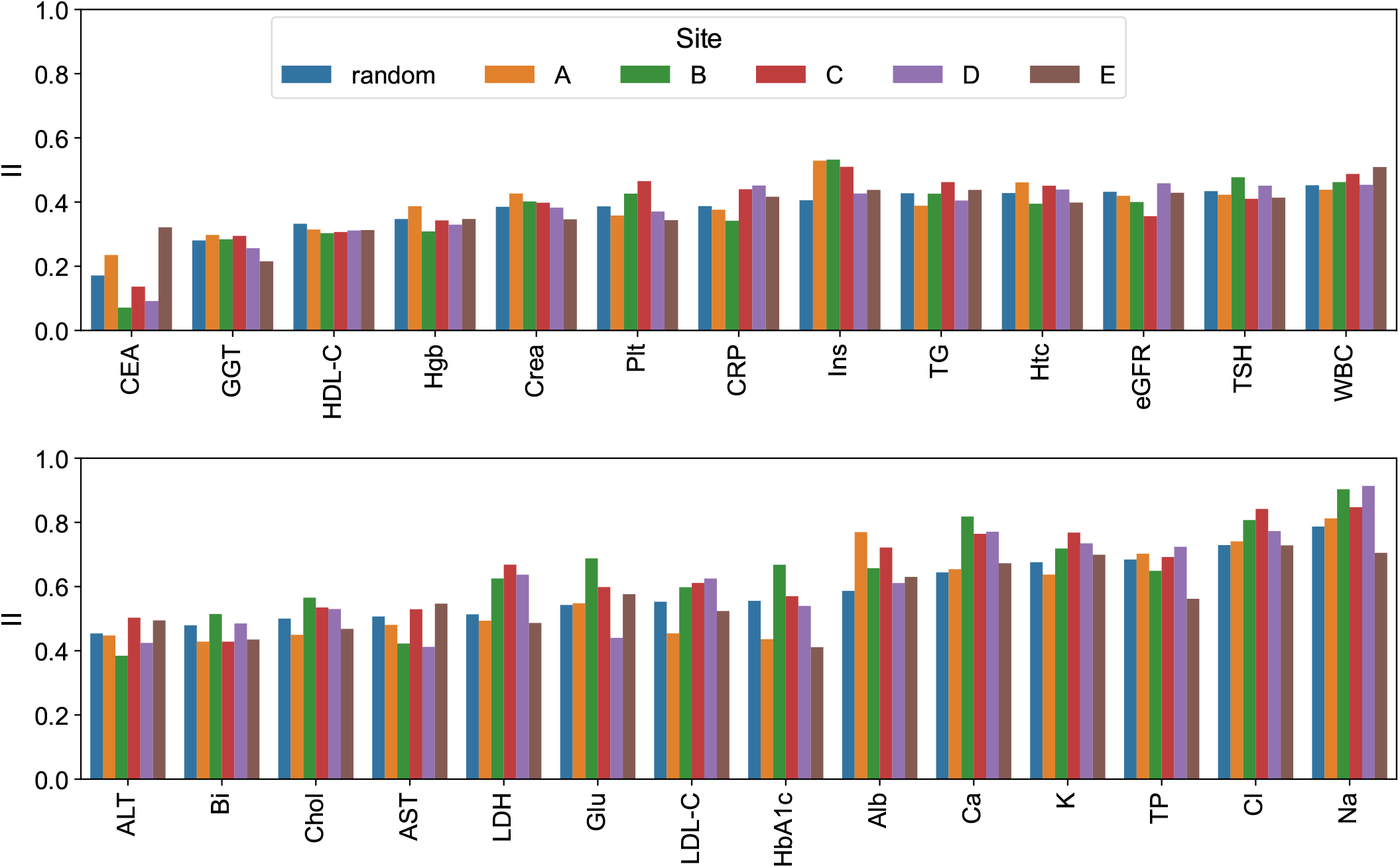
Index of Individuality values for 27 clinical laboratory parameters, calculated in matched subsets of 100 participants each, drawn from the full cohort (random sample) and from each of the five largest study sites (Sites A–E). Participants were matched on age, sex, and BMI.

Additionally, we evaluated whether the molecular fingerprinting performance varied within sites using the same multi-class classification strategy for individual identification as described in the main text (Figure 4b). Learning curves were generated using fixed-size cohorts of 100 randomly selected participants, both from the full dataset and separately from each of the five largest sites. This process was repeated 50 times to estimate the accuracy distributions. Results are shown for FTIR, clinical laboratory, and their combination. The learning curve trends were consistent across all cohorts. Importantly, the combined feature set consistently outperformed the individual modalities, and no substantial drop in accuracy was observed for clinical laboratory models within sites. This suggests that if a strong site-specific bias were present, it would have impaired within-site classification by introducing confounding site effects. However, this was not observed (Figure 14).

**FIG. 14.**
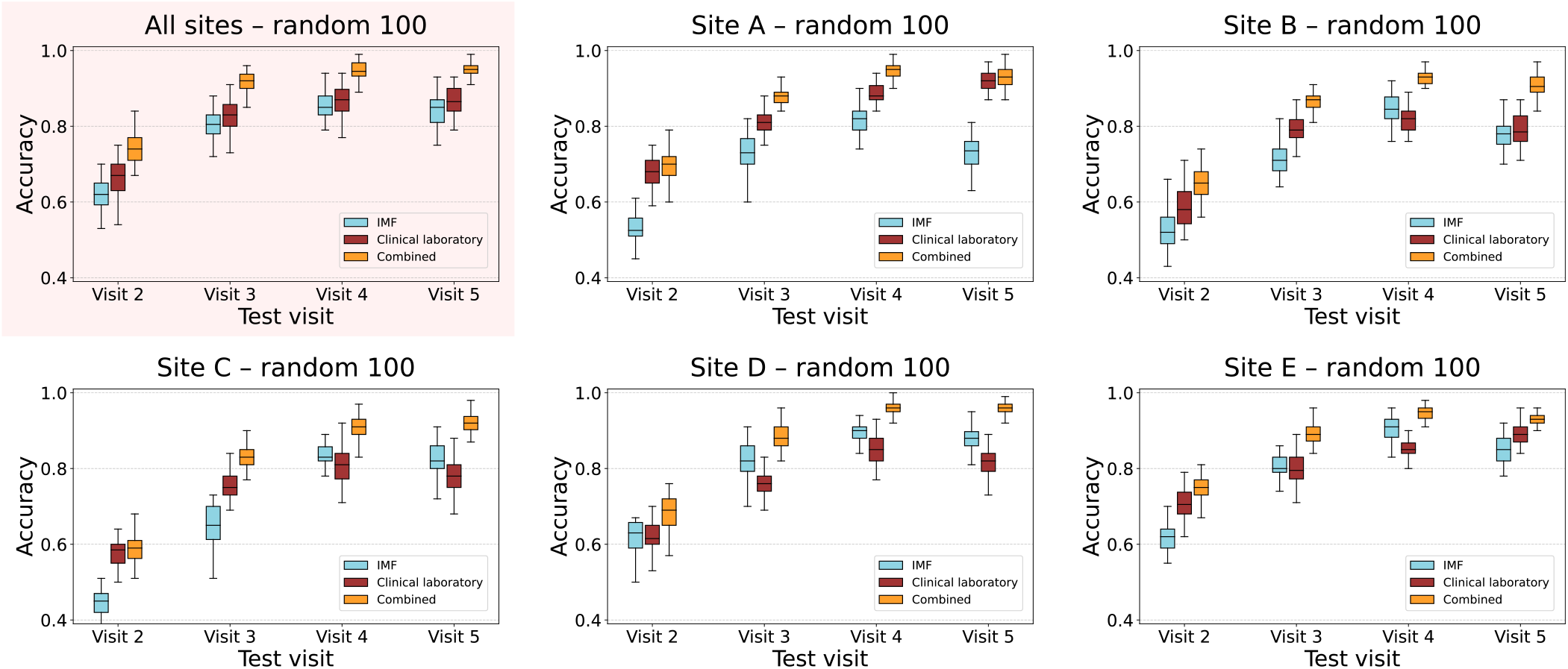
Learning curves for individual identity classification based on FTIR, clinical laboratory, and their combination. Cohorts of 100 participants were randomly sampled from the full dataset and from the five largest sites, with results averaged over 50 repetitions. The consistent performance across sites—particularly for the clinical laboratory modality—supports the robustness of the classification and suggests limited inter-site bias.

## Appendix H: Stability of Infrared Molecular Profiles

To complement the main analysis, we provide a detailed examination of how classification accuracy varies with both the number of training visits and the number of individuals included in the task (Figure 15a). To explore temporal robustness, models were trained on the first *i* visits and evaluated on all subsequent visits to assess re-identifiability based on prior data. For each configuration, we repeated the evaluation 50 times with a randomly selected cohort of size *n*, where *n* ranged from 50 to 3,000, to obtain accuracy distributions.

**FIG. 15.**
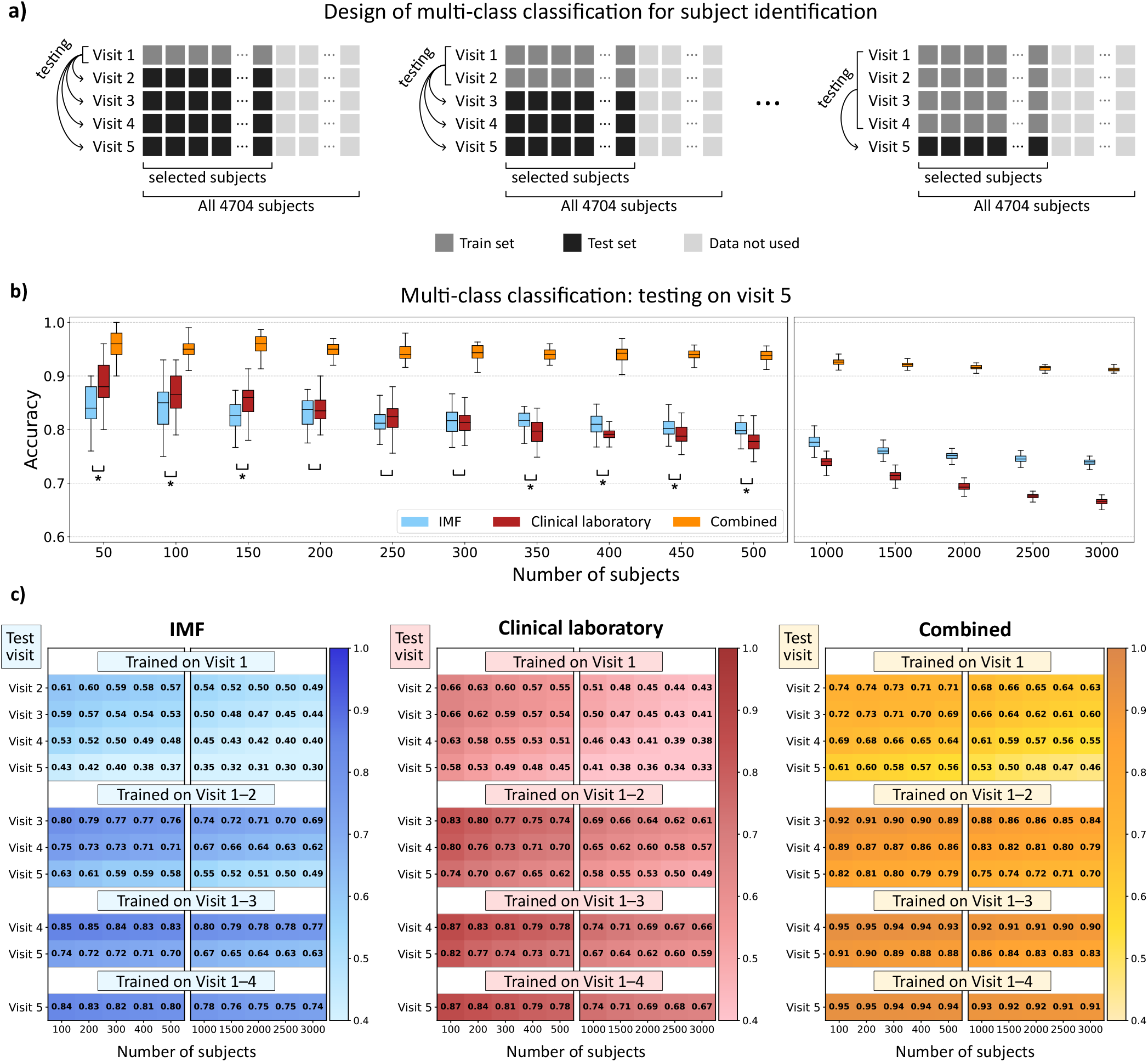
Multi-class classification results for predicting individual identities from longitudinal data across five consecutive visits. Each setup was repeated 50 times with different random subsets to estimate accuracy distributions. **a)** Experimental design: for a randomly selected set of *n* participants, models were trained on the first *i* visits and evaluated on all subsequent visits to assess re-identifiability based on prior data. **b)** Scalability analysis: model performance was assessed across increasing numbers of participants, using the first four visits for training and the fifth for testing, to evaluate accuracy decay with cohort size. Asterisks indicate statistically significant differences (Mann–Whitney U test), with the combined modality consistently outperforming the individual feature sets. As *n* increases, the boxplots become narrower because random subsets increasingly overlap and because each misclassification contributes less to the overall accuracy. For these reasons, boxplots beyond 3,000 participants were not shown, as accuracy distributions could not be meaningfully assessed when sampling from nearly the full cohort. **c)** Temporal generalization: heatmaps show classification accuracy as a function of test visit (rows) and cohort size (columns), for models trained using an increasing number of preceding visits (Visit 1 only, Visit 1–2, Visit 1–3, Visit 1–4). Results are shown for FTIR spectroscopy, clinical laboratory, and the combined feature set. Each cell reflects the mean accuracy from 50 random subsets. Accuracy declined with increasing intervals between training and testing, suggesting a temporal drift in molecular profiles.

When using Visit 5 as the test set and the preceding four visits for training, a clear pattern emerged: clinical laboratory features outperformed IMF in smaller cohorts (fewer than approximately 300 individuals), whereas FTIR surpassed clinical laboratory data at larger scales. Across all cohort sizes, the combined dataset consistently achieved the highest performance (Figure 15b). As the cohort size increased, the boxplots became notably narrower. This is partly because, when drawing increasingly large random subsets from the full cohort, the overlaps between these subsets grow, resulting in less variability between repetitions. Additionally, with very large cohorts, a single additional misclassification affects the overall accuracy by only a very small fraction, which further reduces the spread of accuracy values. For these reasons, beyond 3,000 individuals, the boxplots would have provided little meaningful information, as the accuracy distributions could not be reliably assessed when sampling from the entire pool of 4,704 subjects.

Looking at the overall trends, model performance improved as more visits were included in the training data, and declined as the number of individuals (i.e., classes) increased. Accuracy generally plateaued after the inclusion of four training visits, with slight declines in some settings—most noticeably when using IMF data alone.

To further assess temporal robustness, we evaluated how classification accuracy changed as the temporal gap between training and testing increased (Figure 15c). Models were trained using progressively more prior visits (Visit 1 only, Visits 1–2, Visits 1–3, or Visits 1–4) and tested on later visits. Accuracy decreased with larger temporal gaps, indicating subtle but systematic drift in molecular profiles over time. Nonetheless, the combined feature set mitigated this decline more effectively than either individual modality, again underscoring the benefit of integrating complementary signals.

## Footnotes

1 In the literature, between-person variation is defined as the variability of a central tendency of individuals within a cohort [37]. Under this definition, the index of individuality may exceed 1. In contrast, we define between-subject variation as the average of the individual variabilities across visits. This approach is more natural, as it does not rely on the assumption that a sufficient number of visits per individual are available to estimate average tendencies accurately, and it ensures that the index of individuality remains bounded by 1.

2 It should be emphasized that the definition used here is based on variances, rather than the more common coefficients of variation [38]. This choice is motivated by the fact that average IMF values are often close to zero, which makes the coefficient of variation, in this case, ill-defined.

